# Cultured Bacteria Isolated from Primary Sclerosing Cholangitis Patient Bile Induce Inflammation and Cell Death

**DOI:** 10.1101/2024.10.08.617321

**Authors:** Chelsea E. Powell, Megan D. McCurry, Silvia Fernanda Quevedo, Lindsay Ventura, Kumar Krishnan, Malav Dave, Shaikh Danish Mahmood, Katherine Specht, Raghav Bordia, Daniel S. Pratt, Joshua R. Korzenik, A. Sloan Devlin

**Affiliations:** Department of Biological Chemistry and Molecular Pharmacology, Harvard Medical School, Boston, MA 02115, United States; Division of Gastroenterology, Hepatology and Endoscopy, Brigham & Women’s Hospital, Boston, MA 02115, United States; Autoimmune and Cholestatic Liver Center, Massachusetts General Hospital, Boston, MA 02115, United States

**Keywords:** Primary sclerosing cholangitis, bacteria, microbiome, cellular assays, cellular phenotype

## Abstract

**Background:** Primary sclerosing cholangitis (PSC) is a chronic liver disease characterized by inflammation and progressive fibrosis of the biliary tree. The pathogenesis of PSC remains poorly understood, and there are no effective therapeutic options. Previous studies have observed associations between changes in the colonic and biliary microbiome and PSC. We aimed to determine whether bacterial isolates cultured from PSC patient bile induced disease-associated phenotypes in cells.

**Methods:** Bile was collected from PSC patients (n=10) by endoscopic retrograde cholangiography and from non-PSC controls (n=3) undergoing cholecystectomies. Biliary bacteria were cultured anaerobically, and 50 colonies per sample were identified by 16S rRNA sequencing. The effects of supernatants from seven PSC-associated bacterial strains on cellular phenotypes were characterized using human colonic (Caco-2), hepatic (HepG2), and biliary (EGI-1) cells.

**Results:** No bacteria were isolated from non-PSC controls, while bacteria were cultured from most PSC patients. The PSC bile microbiomes exhibited reduced diversity compared to the gut or oral cavity, with one or two bacterial strains predominating. Overall, PSC-associated bacteria produced factors that were cytotoxic to hepatic and biliary cells. *Enterococcus faecalis*, and to a lesser extent *Veillonella parvula*, induced epithelial permeability, while *Escherichia coli, Fusobacterium necrophorum*, and *Klebsiella pneumoniae* induced inflammatory cytokines in biliary cells.

**Conclusions:** Our data suggest that bacteria cultured from PSC bile induce cellular changes that may contribute to PSC disease pathogenesis. *Enterococcus* may promote intestinal permeability, facilitating bacterial migration to the biliary tree. Once there, *Escherichia, Fusobacterium* and *Klebsiella*, may cause inflammation and damage in biliary and liver cells.

## INTRODUCTION

Primary sclerosing cholangitis (PSC) is a rare, chronic disorder characterized by inflammation and progressive fibrosis of the biliary ducts, which can lead to the development of multifocal biliary strictures, secondary biliary cirrhosis, and decompensated liver disease.^1^ PSC is closely linked with inflammatory bowel disease (IBD); approximately 70-80% of PSC patients have also been diagnosed with IBD.^2,3^ PSC is also associated with a higher risk of cancer in the biliary tract and colon.^1^ Currently there are no effective treatments for PSC other than liver transplant, and PSC can still recur after transplant.^4,5^ The pathophysiology of PSC is poorly understood. Studies into the mechanisms underlying PSC disease phenotypes are essential for the development of new treatments.

While the origins of PSC are unknown, it is currently thought that a combination of environmental and genetic factors contribute to PSC pathogenesis.^2,6^ Among environmental triggers, the gut microbiota is hypothesized to play a key role in PSC development, a theory supported by alterations or dysbiosis detailed in both PSC and IBD.^2^ Previous studies have profiled the gut^7-11^ and biliary^12-14^ microbiome of PSC patients compared to a variety of non-PSC controls. Studies of the gut microbiome in PSC have overall identified lower microbial diversity in PSC patients compared to non-PSC controls, indicative of dysbiosis.^8-11^ Two previous studies of the biliary microbiome identified lower microbial diversity in PSC patients compared to non-PSC controls undergoing endoscopic retrograde cholangiography (ERCP).^12,13^ However, the biliary microbiome is currently understudied and whether or not bile is sterile in healthy livers remains a topic of debate.^15,16^ The gut and biliary microbiomes of PSC patients have been shown to contain an overrepresentation of certain bacterial taxa compared to controls. While the methodologies of these studies were diverse, *Veillonella, Streptococcus, Enterococcus*, and *Fusobacterium* were more abundant in PSC patients consistently across studies.^2,7^ Previous studies have also included fecal^10^ or plasma^9^ metabolomics and observed changes in the abundance of microbiota-derived metabolites in PSC patients compared to controls. Taken together, these alterations in microbial populations and metabolite production suggest that the microbiome may be playing a functional role in the development of PSC. While some studies have used mouse models to explore the therapeutic potential of targeting PSC-derived *Klebsiella pneumoniae* using mouse models,^17,18^ the direct relationship between specific PSC-associated bacteria and disease phenotypes remains underexplored.

Here, we aimed to isolate bacterial strains from PSC patient bile in order to characterize the ability of individual patient isolates to induce disease-associated cellular responses in human colonic, hepatic, and biliary cells. Both our microbial culturing and cellular assays were performed anaerobically to better mimic endogenous conditions. From the 7 PSC-derived strains profiled, we determined that while PSC-associated bacteria in general appeared to produce factors that are cytotoxic to hepatic and biliary cells, other cellular phenotypes were more specific to certain strains. Only *Enterococcus faecalis*, and to a lesser extent *Veillonella parvula*, could induce epithelial permeability, while *Escherichia coli, Fusobacterium necrophorum*, and *Klebsiella pneumoniae* induced the expression of genes encoding inflammatory cytokines in biliary cells. These results suggest that bacteria in PSC patients are causally linked with disease-associated cellular phenotypes and that different species may contribute to distinct aspects of disease pathogenesis.

## METHODS

### Bile Collection

Bile was collected during ERCPs being performed for clinical indications. Informed consent was obtained for a Massachusetts General Hospital/Brigham & Women’s Hospital (Boston, MA, United States) IRB approved protocol (IRB19-1891). Collection methods were designed to minimize contamination from the oral cavity and gastrointestinal tract. Once the scope was introduced into the duodenum the channel was flushed with 20 cc of sterile saline. The catheter was then inserted into the scope and flushed with 5 cc of sterile saline. The catheter was introduced into the biliary tree and at least 3 cc of bile was withdrawn into a sterile empty syringe. The syringe was immediately capped. A second syringe of bile was then drawn until full and immediately capped to limit the amount of headspace in the syringe and minimize potential gas exchange during transport. The first syringe of bile was stored for biobanking and the second syringe was used for bacterial culturing. The sample was not placed on ice or dry ice or otherwise stored, but was immediately transported for culturing. To serve as controls, non-PSC bile was collected during cholecystectomies. Informed consent was obtained. The control bile was obtained soon after cholecystectomy in the operating room. Patients were selected who did not have any active inflammatory disease or cholecystitis and had not been on an antibiotic for at least 4 weeks. A needle was used to pierce through the gallbladder wall and a syringe was used to withdraw at least 3 cc of bile. The syringe was then capped and brought immediately to the lab.

### Bacterial culturing and isolate generation

On the same day of patient bile collection, bacterial culturing was performed in an anaerobic chamber (Coy Laboratory Products) with a gas mixture of 5% H_2_ and 20% CO_2_ (balance N_2_). Pure bile (100 µL) or bile-dilutions in Cullen–Haiser Gut (CHG) media (brain heart infusion (BHI, Bacto) supplemented with 1% BBL vitamin K1-hemin solution (BD), 1% trace minerals solution (ATCC), 1% trace vitamins (ATCC), 5% fetal bovine serum (FBS, Hyclone), 1 g/L cellubiose, 1 g/L maltose, 1g/L fructose, and 0.5 g/L L-cysteine) were plated on CHG agar plates. Dilutions were made at 10^-1^, 10^-2^, 10^-3^, and 10^-4^. Plates were incubated at 37 °C for 2 - 3 days. Individual colonies (50 per sample) were then picked and streaked onto CHG plates for another 2 - 3 day incubation at 37 °C. To ensure isolate purity, the 50 streaked colonies were picked and streaked on CHG plates a second time. After incubation, single colonies from re-streaked agar plates were inoculated into 1 mL of CHG media and incubated at 37 °C for 3 days. An aliquot (10 µL) of each culture was inoculated into 1 mL of SIM+ media (BBL SIM medium (BD) supplemented with 1% BBL vitamin K1-hemin solution (BD) and 0.5 g/L L-cysteine) in order to identify colonies that produced hydrogen sulfide (H_2_S), using *Escherichia coli* Nissle 1917 as a positive control (originally obtained from the Silver laboratory at Harvard Medical School). Another aliquot (500 µL) of the isolate CHG cultures was used for identification by 16S rRNA gene PCR and sanger sequencing (universal 16S-forward, 5’-GAGTTTGATCCTGGCTCAG-3′; universal 16S-reverse, 5′-GGCTACCTTGTTACGACTT-3′), while the remaining isolate cultures were stored at -80 °C as glycerol stocks.

### gDNA extraction from bile and metagenomic sequencing

Genomic DNA (gDNA) was isolated from bile using the PureLink Genomic DNA Mini Kit (Invitrogen) following manufacturer’s instructions for extraction from blood samples. Samples requiring concentration for sequencing were prepared by SeqCenter (Pittsburgh, PA) using AMPure XP Bead-based reagent. Libraries were prepared by SeqCenter using the Illumina DNA Prep Kit (Illumina) and IDT 10 bp UDI indices and sequenced on an Illumina NovaSeq X Plus sequencer, producing 2 x 151 bp paired-end reads resulting in 5 Gbp of data per sample (32M reads). Quality filtering, including demultiplexing and adapter trimming, was performed by SeqCenter using bcl-convert (v4.2.4, Illumina). Taxonomic abund nces were analyzed using MetaPhlan 4.0^19^ and figures were generated using GraphPad Prism 10. Sequencing statistics including total number of reads are reported in Supplemental Table S1. Metagenomic data have been deposited in the NCBI BioProject database (PRJNA1169933) and are publicly available. Accession numbers are listed in Supplemental Table S2.

Attempted isolation of gDNA from bile using the QIAmp DNA Microbiome Kit (Qiagen) or the PowerSoil Pro Extraction Kit (Qiagen), with or without concentration and purification by the PowerClean Pro Kit (Qiagen), did not result in enough gDNA of sufficient purity for metagenomic sequencing. Bile pH was not predictive of extracted gDNA quantity or purity (data not shown).

### Human cell culture

Caco-2 and HepG2 cells were purchased from ATCC, while EGI-1 cells were purchased from DSMZ. Cells were cultured in Dulbecco’s modified eagle media (DMEM) containing glucose and L-glutamine (Gibco), supplemented with 10% FBS (Genclone) and 1% penicillin/streptomycin. Mycoplasma testing was performed using a MycoAlert Mycoplasma Detection Kit (Lonza) and all lines were negative.

### Bacterial supernatant collection for cellular assays

Bacterial cultures were diluted to an OD_600_ of 0.1 in CHG media and grown anaerobically at 37 °C overnight before the day of human cell assays. After incubation, bacterial supernatants were collected inside the anaerobic chamber by sterile-filtration using a 0.2 µm Supor Membrane syringe filter (Pall Corporation) in order to maintain any bacterially produced gases as previously described.^20^ Collected supernatants were immediately used in cell assays. *Enterococcus gallinarum* (gift from Martin Kriegel at Yale School of Medicine)^21^ and *Bacteroides fragilis* (ATCC 25285) were used as reference strains and were not derived from PSC patients.

### Cell viability

The day before treatment, 30,000 Caco-2, HepG2, or EGI-1 cells were plated in 100 µL of complete DMEM media in a 96-well white Nunclon Delta-treated plate (ThermoFisher) and incubated at 37 °C with 5% CO_2_ overnight. The day of the assay, media was aspirated and cells were moved into the anaerobic chamber. Anaerobic complete DMEM media was added to the wells and cells were incubated at 37 °C for 1 h before adding sterile-filtered bacterial supernatants such that the final well volumes were 100 µL of supernatant and DMEM at ratios of 1:1 or 1:3 (v/v). Cells were then incubated anaerobically at 37 °C for 16 h before viability was assessed using the CellTiter-Glo luminescent cell viability assay (Promega), following manufacturer’s standards. Luminescence signal was normalized to wells with supernatant and DMEM without human cells.

### Epithelial permeability assay

Caco-2 cells were differentiated in 24-well plate transwells (0.4 µM pore size, Costar) as previously described.^22^ On the day of the assay, differentiated Caco-2 cells were washed with Hank’s balanced salt solution (HBSS, Gibco) before being moved into the anaerobic chamber. Anaerobic HBSS (500 µL) was added to the basal chamber and 50 µL of anaerobic HBSS with 10 µM 4 kDa FITC-Dextran (Sigma-Aldrich) was added to each apical chamber before incubation at 37 °C for 1 h. Syringe-filtered bacterial supernatants (50 µL) were then added to each well and cells were incubated at 37 °C for 6 or 12 h. HBSS (100 µL) from the basal chamber was then moved to a black 96-well plate (Corning) and fluorescence was measured using an Envision 2104 plate reader at Harvard Medical School’s ICCB-Longwood Screening Facility.

### Gene expression analysis by qRT-PCR

The day before treatment, 1,000,000 EGI-1 or HepG2 cells were plated in 2 mL complete DMEM media in 6-well Nunclon Delta-treated plates (ThermoFisher) and incubated at 37 °C with 5% CO_2_ overnight. The day of the assay, media was aspirated and cells were moved into the anaerobic chamber.

Anaerobic complete DMEM media (1 mL) was added to the wells and cells were incubated at 37 °C for 1 h. Next, sterile-filtered bacterial supernatant (1 mL) was added to each well and cells were incubated at 37 °C for 6 h. Cells were then removed from the anaerobic chamber, media was aspirated, and cells were washed with 1 mL ice cold PBS. Cells were lysed in ice cold TRI Reagent (Zymo Research) and stored at -80 °C until RNA was extracted using the Direct-zol RNA Miniprep Plus kit (Zymo Research) with DNase cleanup step. RNA (1 µg per sample) was reverse transcribed using High-Capacity cDNA Reverse Transcription kit (Applied Biosystems). cDNA (10 ng per sample) was then analyzed by qRT-PCR using LightCycler 480 SYBR Green I Master (Roche). Reactions were performed in a 384-well format on a QuantStudio 7 Pro at Harvard Medical School’s ICCB-Longwood Screening Facility. The 2^-ΔΔCt^ method^23^ was used to calculate relative changes in gene expression and all results were normalized to UBC mRNA expression. Primer sequences were obtained from PrimerBank (https://pga.mgh.harvard.edu/primerbank/). Primer efficiencies were determined to be between 90 – 110% with R^2^ values between 0.99 and 1.00. Sequences are provided in the Supplementary Information (Supplemental Table S3).

## RESULTS

### PSC patient bile contains anaerobically culturable bacteria

We collected bile from 10 individuals with PSC by ERCP as well as from 3 individuals undergoing cholecystectomies without PSC who served as control subjects (Table 1, Supplemental Table S4). Anaerobic bacterial culturing of these samples demonstrated that the majority of the PSC patient bile samples contained culturable bacteria. In contrast, none of the control samples contained culturable microbes (Table 1, Figure 1). Colony picking followed by 16S rRNA sequencing of restreaked isolates further revealed that most of the bile samples of PSC patients were dominated by 1 - 2 strains of bacteria (Figure 1). Two bile samples taken from the same patient 8 months apart demonstrated that it is possible for the same 1 - 2 strains to dominate the biliary microbiome in the same individual over time (Figure 1B). These isolates were also examined for their ability to produce H_2_S, which has been linked to intestinal dysbiosis and IBD, particularly ulcerative colitis (UC).^24-28^ Of the PSC samples with culturable bacteria, half (4 of 8) contained H_2_S-producing strains (Supplemental Tables S5 – S13). Metagenomic analysis, while not possible for all samples due to the difficulty of gDNA extraction from bile (see Methods), generally validated our PSC bile culturing results. Most samples exhibited a similar overabundance of the same 1 - 2 strains of bacteria that we observed in our bacterial culturing (Supplemental Figure S1). However, metagenomic analysis of bile samples from PSC 4 and PSC 9 did reveal the presence of several unculturable species. Overall, these results show that there were live bacteria in the bile from the majority of PSC patients in this cohort. While most samples contained one or two dominant strains, the identity of these strains differed from patient to patient, highlighting the heterogeneity of individual PSC biliary microbiomes.

**Table 1.** Patient characteristics at time of bile collection.

|  | **PSC** | **Controls** |
| --- | --- | --- |
| **Patients, n** | 10 | 3 |
| **Age (years), median (range)** | 39.5 (28-74) | 41 (35-59) |
| **Female, n (%)** | 2 (20) | 2 (66.7) |
| **Age at Diagnosis (years), median (range)** | 34.5 (23-70) | NA |
| **Disease Duration (years), median (range)** | 8 (0-28) | NA |
| **IBD, n (%)** | 8 (80) | 0 |
| **Age at IBD Diagnosis (years), median (range)** | 26 (16-48) | NA |
| **IBD Duration (years), median (range)** | 14 (3-55) | NA |
| **Other Inflammatory Conditions, n (%)** | 1 (10) | 1 (33.3) |
| **ERCP (#), median (range)** | 4 (1-8) | 0 |
| **ALP (U/L), median (range)** | 380.5 (58-629) | 86 (41-106) |
| **Antibiotic Use 3 Months Prior to Collection, n (%)** | 10 (100) | 0 |
| **Corticosteroids, n (%)** | 2 (20) | 0 |
| **5-ASA, n (%)** | 4 (40) | 0 |
| **Biologics, n (%)** | 2 (20) | 0 |
| **UCDA, n (%)** | 6 (60) | 0 |
| **Proton Pump Inhibitors, n (%)** | 1 (10) | 0 |
| **Culturable Bacteria Species, median (range)** | 2 (0-23) | 0 (0-0) |

**Figure 1.**
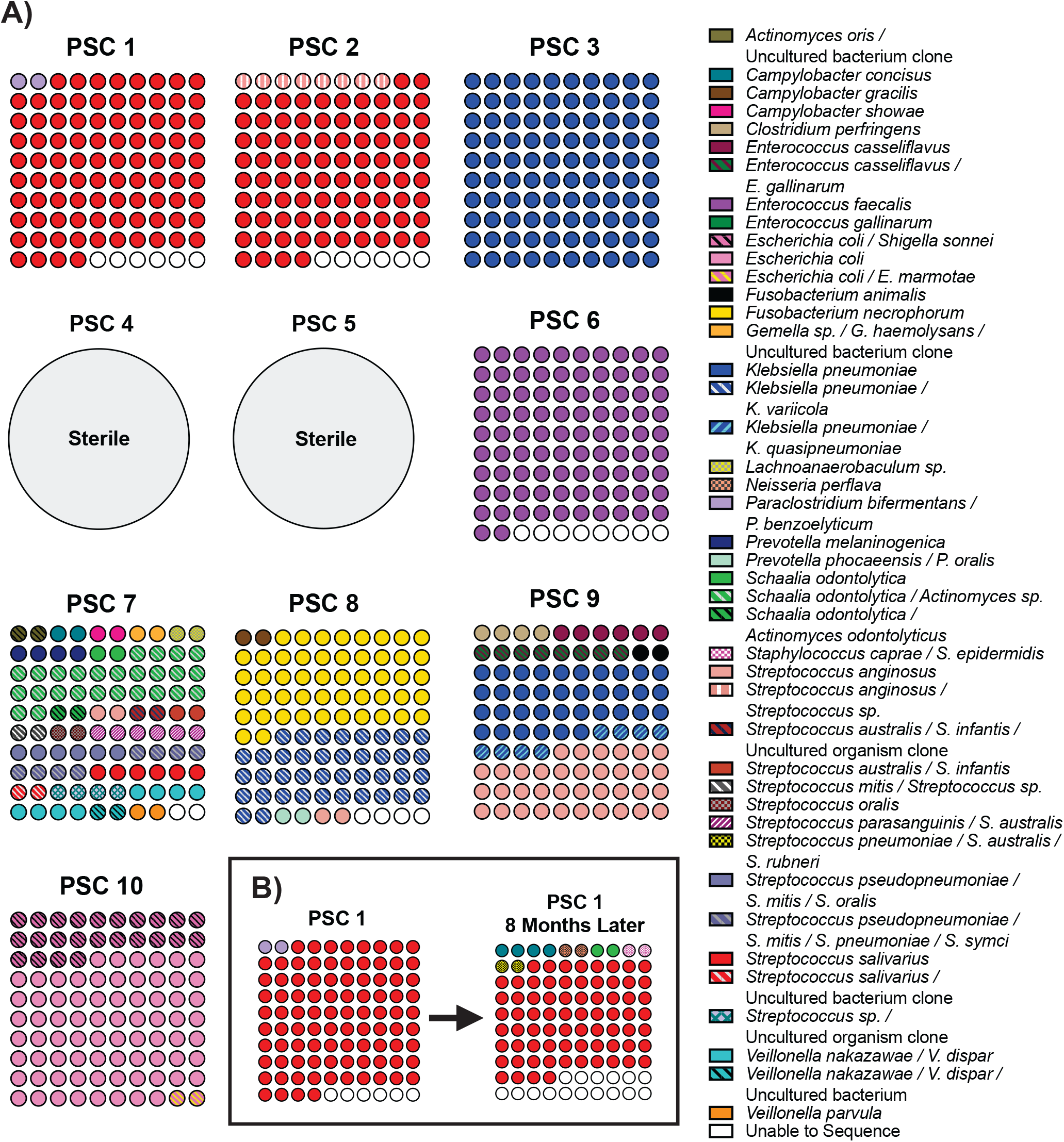
PSC patient bile contains culturable bacterial isolates. A) Fifty bacterial colonies were isolated from each distinct patient bile sample under anaerobic conditions. Bacteria were identified by 16S rRNA gene PCR and Sanger sequencing. 〇〇 = 1 colony out of 50 (2%). B) Anaerobically culturable bacteria were isolated from bile samples from the same patient, 8 months apart. For complete lists of cultured bacteria in each sample, see Tables S5-S13.

### PSC bacterial isolates produce factors that affect human cell viability

Seven bacterial isolates generated from the PSC bile culturing were selected for further characterization using human cellular assays (Table S14). These strains were selected due to their abundance in PSC patient bile in our study (*Enterococcus faecalis, Escherichia coli, Fusobacterium necrophorum, Klebsiella pneumoniae*, and *Streptococcus salivarius*) or because they were identified as abundant in PSC patient microbiomes in previous studies (*Veillonella dispar/nakazawae* and *Veillonella parvula*).^8,10,11,29,30^ *Bacteroides fragilis* ATCC 25285,^31^ a human gut commensal strain that has been shown to promote epithelial integrity, and *Enterococcus gallinarum* (isolated by Kriegel lab at Yale School of Medicine), a pathogenic strain that has been shown to induce intestinal permeability and translocate from the gut to the liver,^21^ were selected for comparison to the PSC-derived isolates. Importantly, *B. fragilis* ATCC 25285 is an enterotoxin-free strain that has been characterized as a protective probiotic^31,32^ and is distinct from the subset of enterotoxigenic *B. fragilis* associated with UC.^33,34^

Cell viability assays were performed using established colonic (Caco-2), biliary (EGI-1), and hepatic (HepG2) human cancer cell lines under anaerobic conditions. The effects on cell viability of gut commensal reference strain *B. fragilis* and pathogenic reference strain *E. gallinarum* were otherwise previously unknown against these specific human cell lines. The gut commensal control, *B. fragilis*, did not induce cell death in the colonic cell line (Figure 2).^31^ *B. fragilis* instead induced growth in Caco-2 cells, and induction of slight growth has indeed been seen in a previous study using normal colonic epithelial cells (hcoEPIC).^31^ *E. gallinarum* induced cell death in all three cell lines, while *B. fragilis* induced cell death in the biliary and hepatic cell lines. The fact that *B. fragilis* is gut commensal that does not usually interact with biliary and hepatic cells may have contributed to its cytotoxic effects on these cells types. Sterile-filtered bacterial supernatants from all but one bacterial isolate induced cell death in both biliary and hepatic cells, with hepatic cells being affected to a greater extent (Figure 2). Notably, the PSC bacterial supernatants, with the exception of *E. faecalis* and *F. necrophorum*, induced cell growth instead of cell death in colonic cells. *V. parvula* also induced growth in hepatic cells and, to a lesser extent, in biliary cells. These results indicate that some biliary bacteria are producing toxic factors that negatively impact viability in biliary and hepatic cells, but not colonic cells, suggesting that these bacterial factors exert cell-specific effects.

**Figure 2.**
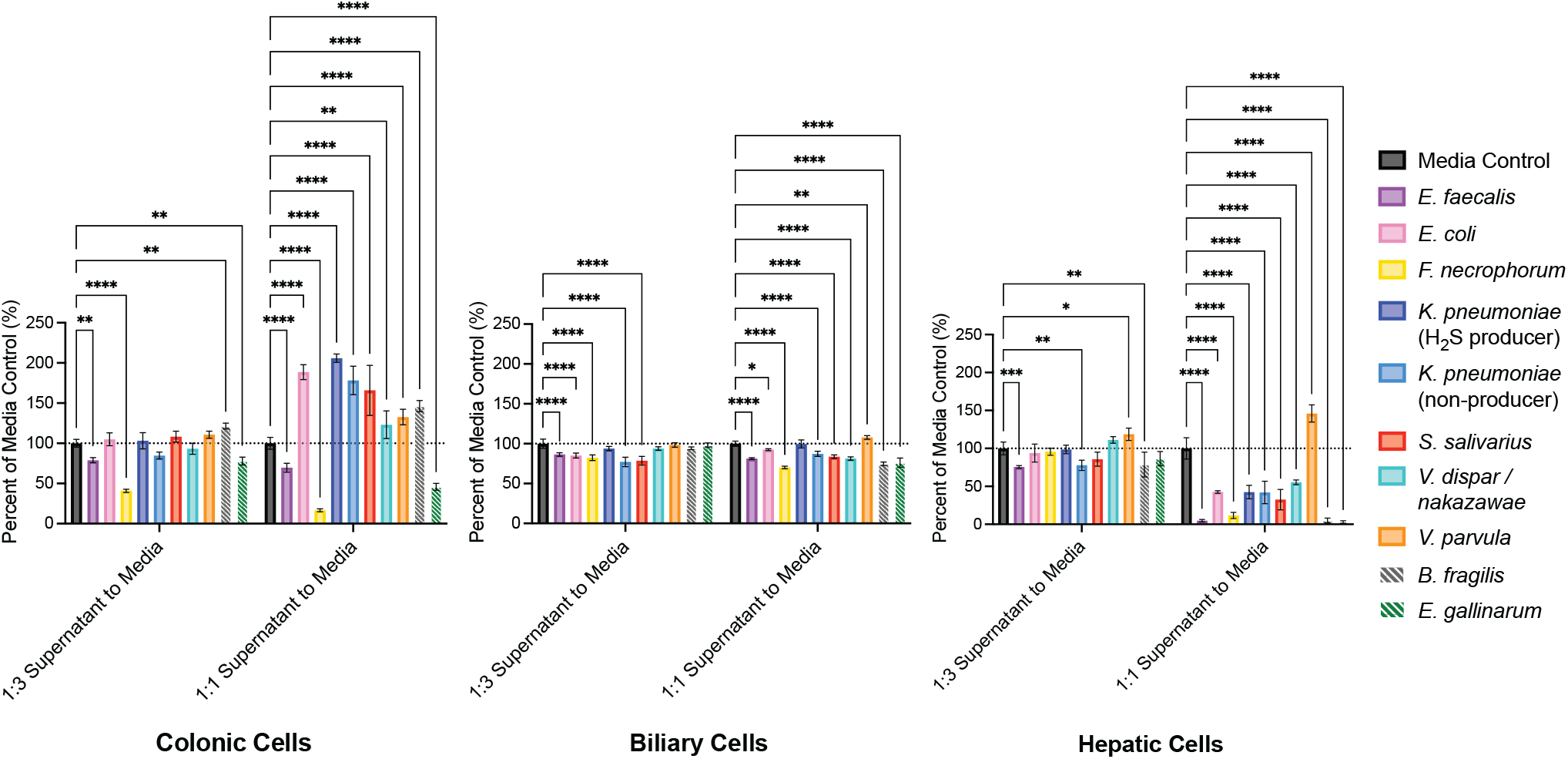
PSC bacterial supernatants affect human cell viability. Colonic (Caco-2), biliary (EGI-1), and hepatic (HepG2) human cancer cells were treated with a 1:3 or 1:1 ratio (v/v) of bacterial culture supernatant for 16 h under anaerobic conditions. CHG media was used as a control. *B. fragilis* and *E. gallinarum* were used as reference strains and were not derived from PSC patients, as indicated by striped bars. Two-way ANOVA was performed followed by Dunnett’s multiple comparisons test using Graphpad Prism 10 software (values are shown as mean ± SD; four biological replicates; \**p* < 0.05, \*\**p* < 0.01, \*\*\**p* < 0.001, \*\*\*\**p* < 0.0001).

### PSC patient-derived *E. faecalis* and *V. parvula* induce intestinal permeability

We used a Caco-2 transwell assay to model induction of pathogenic intestinal permeability (also known as “leaky gut”).^22,35^ Reference strain *B. fragilis* did not induce permeability, while reference strain *E. gallinarum* caused barrier damage, as expected from previous studies.^21,31,36^ Of all the PSC-associated bacteria examined, only supernatant derived from *E. faecalis* cultures induced epithelial permeability after both 6 and 12 h of treatment. *V. parvula* supernatant induced permeability at 12 h alone (Figure 3). Taken together with the colonic cell viability data, these results suggest that *E. faecalis* may be contributing to both gut epithelial cell death and barrier damage, while an array of PSC strains may be causing cell damage in the biliary tree and liver.

**Figure 3.**
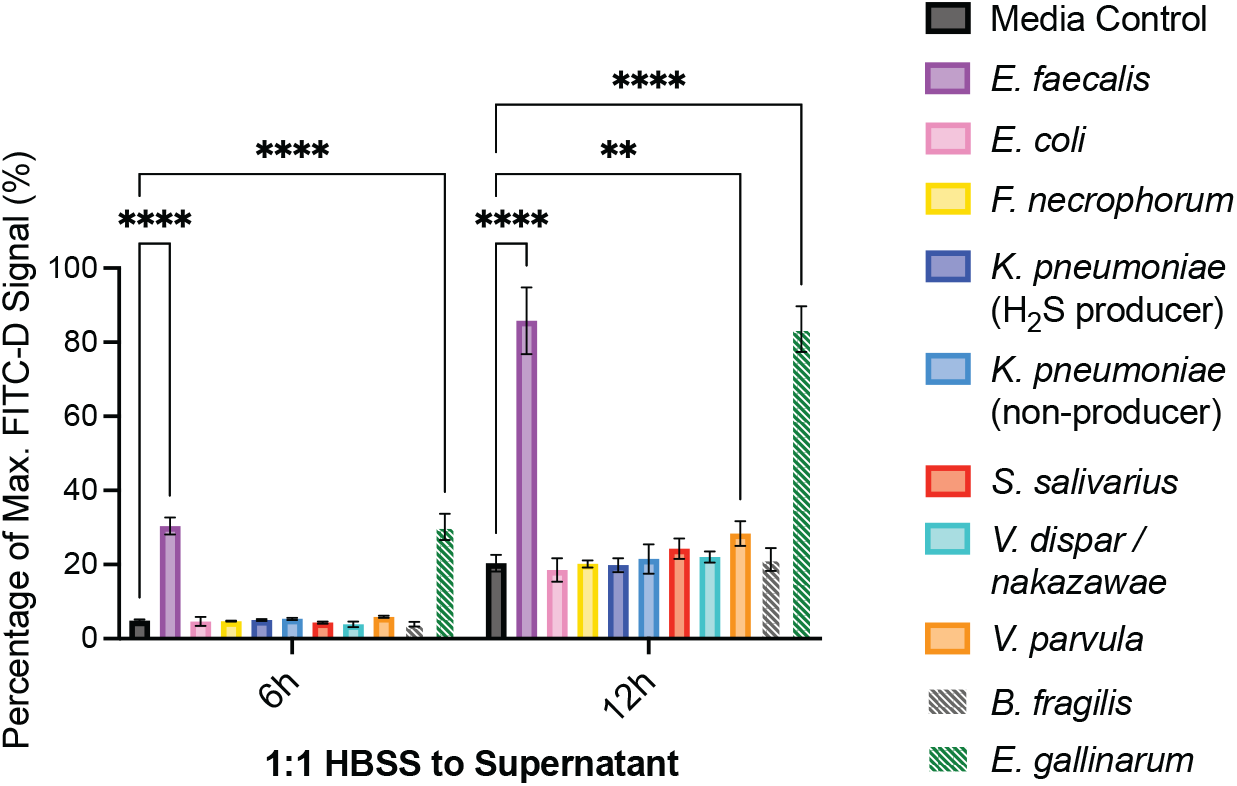
Bacterial supernatants from the PSC patient isolates *E. faecalis* and *V. parvula* increased epithelial permeability. A differentiated monolayer of Caco-2 cells was treated with a 1:1 ratio (v/v) of HBSS buffer to bacterial culture supernatant for 6 or 12h under anaerobic conditions. CHG media was used as a control. *B. fragilis* and *E. gallinarum* were used as reference strains and were not derived from PSC patients, as indicated by striped bars. Two-way ANOVA was performed followed by Dunnett’s multiple comparisons test using Graphpad Prism 10 software (values are shown as mean ± SD; four biological replicates; \**p* < 0.05, \*\**p* < 0.01, \*\*\**p* < 0.001, \*\*\*\**p* < 0.0001).

### PSC bacterial isolates induce inflammatory cytokines in biliary cells

qPCR after 6 h treatment with PSC-derived bacterial supernatant indicated that *E. coli, F. necrophorum*, and *K. pneumoniae* induced transcription of the pro-inflammatory cytokines TNFα and IL-8 in biliary cells but did not exert the same effect in hepatic cells. *V. parvula* induced inflammatory responses in both cell types (Figure 4). Our results show that a subset of PSC-associated bacteria induce inflammatory responses in biliary cells and identify TNFα and IL-8 as the primary cytokines induced by these strains.

**Figure 4.**
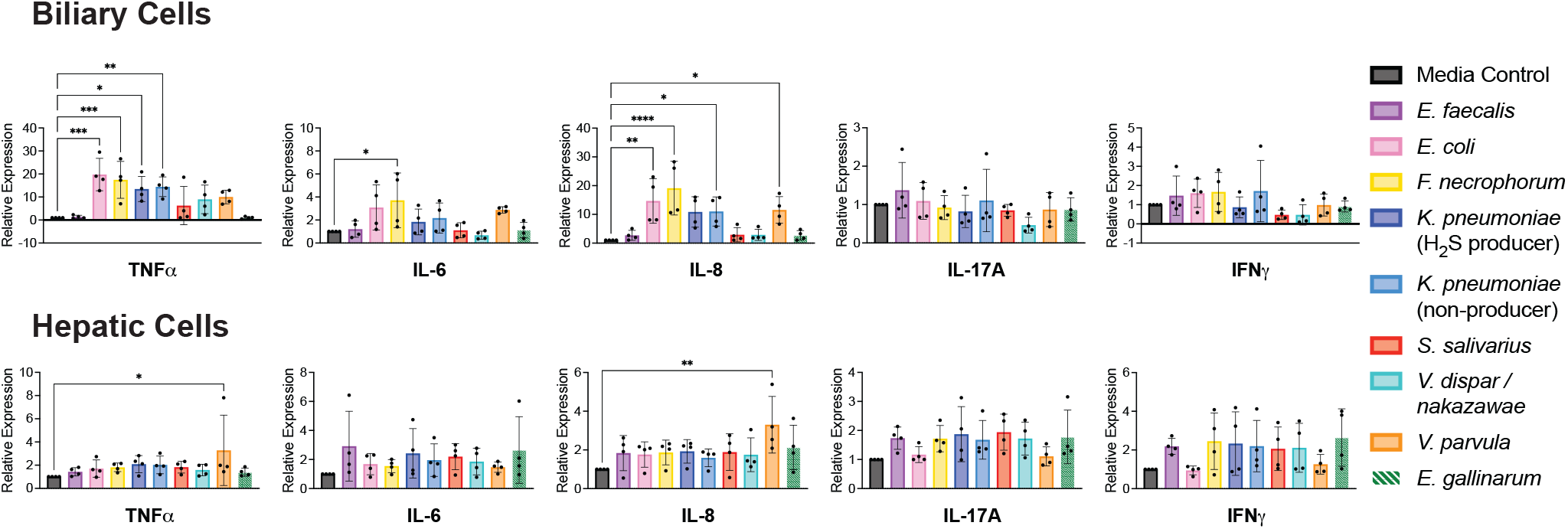
PSC bacterial supernatants induced transcription of inflammatory cytokines in biliary and hepatic cells. Biliary (EGI-1) and hepatic (HepG2) cells were treated with a 1:1 ratio (v/v) of bacterial culture supernatant to cell culture media for 6 h under anaerobic conditions and effect on transcription of target genes was assessed by qPCR. CHG media was used as a control. *E. gallinarum* was used as a reference strain and was not derived from PSC patients, as indicated by striped bars. Results were normalized to UBC mRNA expression. One-way ANOVA was performed followed by Dunnett’s multiple comparisons test using Graphpad Prism 10 software (values are shown as mean ± SD; four biological replicates from independent experiments with three technical replicates each; \**p* < 0.05, \*\**p* < 0.01, \*\*\**p* < 0.001, \*\*\*\**p* < 0.0001).

### PSC-derived bacteria can affect protective pathways in human biliary and hepatic cells

Next, expression of targets involved in sulfur metabolism, mucin expression, and the bicarbonate umbrella were measured by qPCR in order to assess whether PSC-associated bacteria affected the ability of biliary and hepatic cells to protect themselves from damage. Supernatants from *V. dispar/nakazawae* cultures caused a decrease in mRNA expression of sulfide quinone oxidoreductase (SQOR) in biliary cells, which indicates damage to a pathway that aids in the clearance of toxic sulfides (Figure 5). *S. salivarius* and *E. faecalis* supernatants upregulated expression of sulfur metabolism targets in both hepatic and biliary cells, as well as expression of AE2, a primary component of the protective bicarbonate umbrella in biliary cells. *S. salivarius* also upregulated mucin expression in hepatic cells, while *V. parvula* upregulated mucin expression in biliary cells. Increased expression of genes involved in sulfur metabolism, mucin expression, and the bicarbonate umbrella may indicate that hepatic and biliary cells initially upregulate protective pathways in response to toxic factors produced by these bacteria.

**Figure 5.**
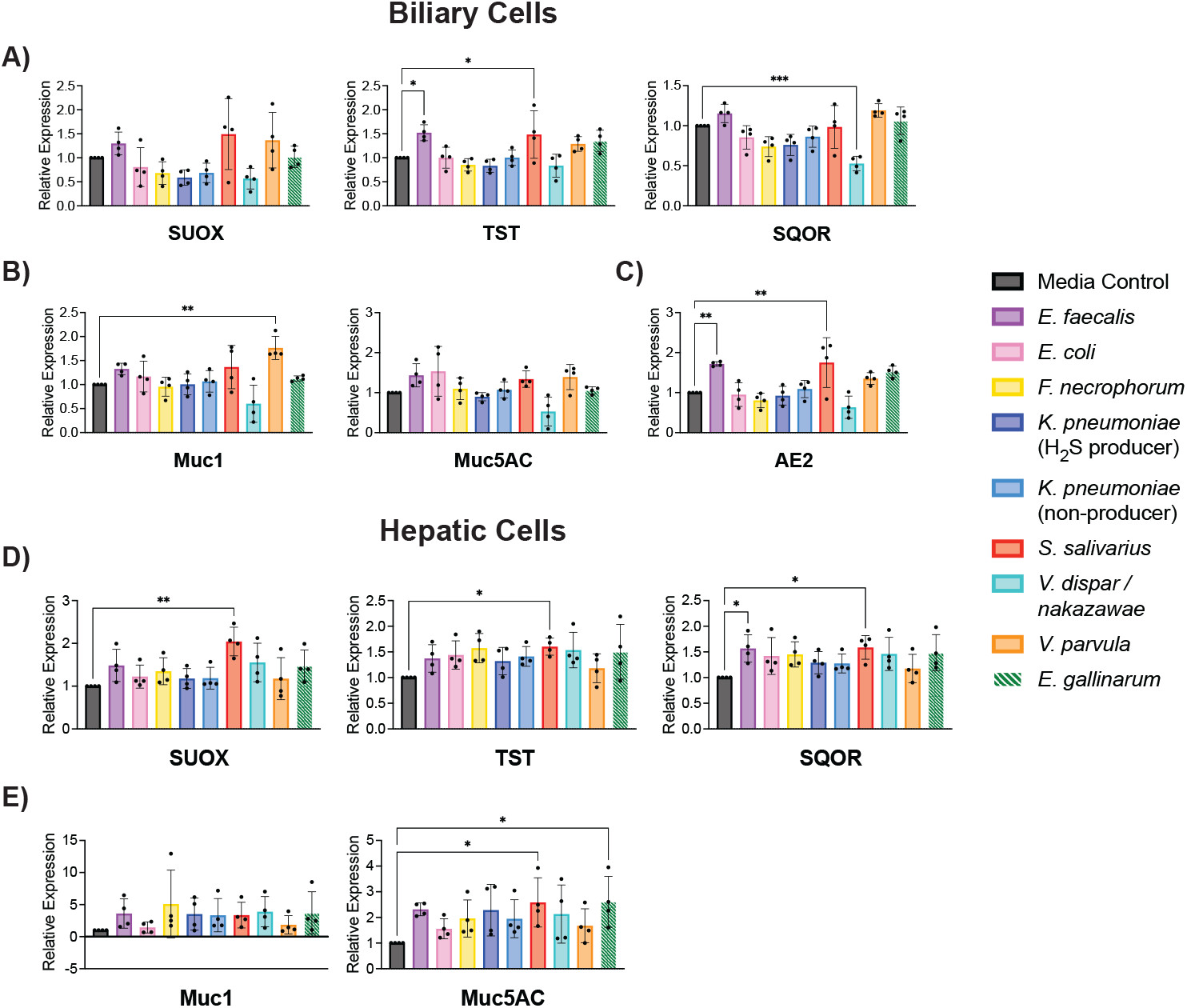
PSC bacterial supernatants affected transcription of genes related to sulfur metabolism, mucin expression, and the bicarbonate umbrella in biliary and hepatic cells. Biliary (EGI-1) and hepatic (HepG2) cells were treated with a 1:1 ratio (v/v) of bacterial culture supernatant to cell culture media for 6 h under anaerobic conditions and effect on transcription of target genes was assessed by qPCR. CHG media was used as a control. *E. gallinarum* was used as a reference strain and was not derived from PSC patients, as indicated by striped bars. Results were normalized to UBC mRNA expression. One-way ANOVA was performed followed by Dunnett’s multiple comparisons test using Graphpad Prism 10 software (values are shown as mean ± SD; four biological replicates from independent experiments with three technical replicates each; \**p* < 0.05, \*\**p* < 0.01, \*\*\**p* < 0.001, \*\*\*\**p* < 0.0001).

## DISCUSSION

The pathogenesis of PSC remains poorly understood and examining potential mechanistic drivers of the disease is essential for developing new treatments. Previous studies have established that PSC patients have altered gut and biliary microbiomes compared to non-PSC control subjects.^7-14^ Here, we sought to identify causal links between the PSC microbiome and disease-associated cellular phenotypes. Through the use of PSC patient-derived bacterial isolates and human cellular assays, we have identified bacterial species that induce PSC-associated cellular phenotypes, including epithelial permeability, biliary and hepatic inflammation, and cell death (Table 2). Importantly, our bacterial and cell culture experiments were performed under anaerobic conditions, allowing us to better replicate the endogenous conditions in the biliary tree and GI tract. Interestingly, from the strains examined we observed that different species induce distinct cellular responses. Our results suggest that the overall microbial community in PSC patients contributes to disease development as opposed to single strains acting as universal drivers of pathogenesis.

**Table 2.** Summary of characterization of bacteria isolated from PSC patient bile.

|  | Description | Gram +/- | H<sub>2</sub>S Producer? | Biliary Viability | Hepatic Viability | Colonic Viability | Intestinal Permeability | Biliary Inflammation | Hepatic Inflammation |
| --- | --- | --- | --- | --- | --- | --- | --- | --- | --- |
| *Enterococcus faecalis* | GI tract commensal | + | No | Killing | Killing | Killing | Induces | No effect | No effect |
| *Escherichia coli* | GI tract commensal | - | Yes | Killing | Killing | Induces growth | No effect | Induces | No effect |
| *Fusobacterium necrophorum* | Throat pathogen | - | Yes | Killing | Killing | Killing | No effect | Induces | No effect |
| *Klebsiella pneumoniae* | Skin, mouth and GI tract commensal | - | Mixed | Killing | Killing | Induces growth | No effect | Induces | No effect |
| *Streptococcus salivarius* | Oral and upper respiratory commensal | + | No | Killing | Killing | Induces growth | No effect | No effect | No effect |
| *Veillonella dispar/ nakazawae* | Oral commensals | - | No | Killing | Killing | Induces growth | No effect | No effect | No effect |
| *Veillonella parvula* | Oral commensal | - | No | Induces growth | Induces growth | Induces growth | Induces | Induces | Induces |

First, we observed that in most patients, one or two bacterial strains predominated in individual PSC patient bile. This finding agrees with previous studies that have demonstrated that the PSC gut and biliary microbiomes contain an overabundance of certain species compared to non-PSC controls.^2,12^ Culturing of our non-PSC controls requiring cholecystectomies did not yield any bacterial isolates. In contrast, two previous PSC biliary microbiome studies that found greater microbial diversity in non-PSC compared to PSC bile.^12,13^ However, the non-PSC controls in these previous studies were from patients without PSC that required an ERCP, not cholecystectomy patients. Comparing our results with this previous work highlights the need to further characterize the biliary microbiome in different disease states. Our non-PSC cholecystectomy bile results align more closely with a recent study which found that bile from pathologically normal gallbladders did not contain culturable microbes.^37^ In contrast, *Enterococcus faecalis, Escherichia coli, Fusobacterium necrophorum, Klebsiella pneumoniae, Streptococcus anginosus*, and *Streptococcus salivarius* were abundant cultured strains in PSC bile (Figure 1). One limitation of our study is the relatively small number of patients samples collected. Despite this limitation, our results are consistent with previous PSC microbiome profiling efforts,^7^ which observed an overabundance of *Enterococcus, Fusobacterium*, and *Streptococcus* in PSC patients. Thus, it is likely that our samples are representative of at least a substantial subset of PSC patient microbiomes that have been consistently identified in the literature with the exception that we did not see an overabundance of *Veillonella*, as has been observed in previous studies.^8,10,11,29,30^. Importantly, our culturing results demonstrate that live bacteria are present in the bile of PSC patients, a conclusion that is not possible based on sequencing-based analyses alone.

In order to identify non-culturable bacteria in the bile strains, we performed metagenomic sequencing. However, despite multiple attempts using different techniques (see Methods), gDNA proved difficult to extract from bile, preventing metagenomic analysis of some patient bile samples. Overall, the bacterial identity and diversity of each sample that we observed in our metagenomic analyses were similar to those from our culturing data. These results suggest that the lower abundance of *Veillonella* in our isolated strains compared to previous PSC microbiome studies was not a result of our culturing methods but indeed reflective of our patient population. For two of the sequenced samples (PSC 4 and PSC 9), we identified some bacteria that we were unable to culture, including strains from the genera *Sphingomonas, Acidovorax*, and *Campylobacter* (Supplemental Figure S1). The identification of unculturable bacteria is of particular note for PSC 4, which appeared sterile during our culturing efforts, suggesting that some bacterial species were not amenable to our culturing methods. No bacteria were able to be identified by metagenomic analysis in PSC 7 and PSC 1 (8 months later), despite successful culturing in these samples. One possible explanation for the failure of sequencing in these cases is that gDNA extraction resulted in the collection of mainly eukaryotic DNA, and a negligible amount of bacterial DNA was isolated. This was indeed seen with PSC 7 and PSC 1 (8 months later) where there was only 0.4% and 1% bacterial DNA relative abundance, respectively, seen by profiling using Kraken2^38^ followed by KronaTools^39^ (Supplemental Table S15). Our work highlights the difficulty of extracting bacterial DNA from some bile samples and motivates the development of more robust DNA isolation methods given the heterogeneity of bile sample characteristics, including viscosity and composition.

In cell culture assays, supernatants from bacteria that we had isolated from PSC bile induced cell death in both biliary and hepatic cells, with hepatic cells being affected to a greater extent (Figure 2).

These results may suggest that cells in the biliary tree can tolerate these bacteria, despite the fact they are producing cytotoxic factors. This resistance to cell death could potentially allow for bacteria to colonize the biliary tree, leading to a buildup of these cytotoxic factors that can then cause liver damage. Interestingly, the PSC bacterial supernatants, with the exception of *E. faecalis* and *F. necrophorum*, induced cell growth in the colonic cancer cell line. These results may indicate that these bacteria produce prooncogenic factors, a finding that is consistent with the association of PSC with increased risk of colorectal cancer.^1^ *V. parvula* was unique among the PSC-derived bacteria in that it did not induce cell death, but instead induced growth in hepatic cells and, to a lesser extent, in biliary cells. Indeed, an increased abundance of *V. parvula* has been correlated with intrahepatic cholangiocarcinoma.^40^ Future work is needed to identify potential prooncogenic factors produced by PSC-associated bacterial strains. While human cancer cell lines are established as useful *in vitro* models due to ease of cell culturing, it should be noted that they may be more responsive, particularly to prooncogenic factors, by nature of being cancer-derived and already in a disease state. Therefore, the effects of these bacteria on colonic, hepatic, and biliary cell health in PSC patients may be more subtle. Of note, the reference commensal strain *B. fragilis* (ATCC 25285), which was not derived from PSC patients, induced growth in colonic cells and cell death in hepatic and biliary cells, despite being an enterotoxin-free strain that is generally considered protective and non-pathogenic (Figure 2).^31,32,36,41,42^ It is possible that these results with *B. fragilis* are reflective of an increased responsiveness in these cancer cell lines.

Our intestinal permeability assay results demonstrate that *E. faecalis* is a strong inducer of epithelial barrier damage and *V. parvula* is a weaker inducer, while other PSC-derived bacteria did not have this effect (Figure 3). These data suggest that *Enterococcus* may be a driver of pathogenic intestinal permeability in PSC patients, potentially allowing for gut bacteria to translocate from the gut to the liver via the portal vein and colonize the biliary tree. These results appear consistent with previous studies that have identified an increase in abundance of *E. faecalis* in PSC ductal bile compared to controls without PSC.^12^ While *E. faecalis* is present in human gut microbiomes, including those of PSC patients,^30,43^ we do not have paired fecal samples from the PSC patients in this study to confirm whether individuals with *E. faecalis* in their bile also have this microbe in their GI tract. Future studies pairing gut and bile metagenomic sequencing will help reveal connections between bacteria that induce intestinal permeability and biliary tree colonization. Additionally, future studies may benefit from examining if *E. faecalis* induces biliary permeability in mouse models or in *in vitro* models of biliary epithelial permeability.

We identified the PSC-associated bacteria *E. coli, F. necrophorum, K. pneumoniae*, and *V. parvula* as inducers of inflammatory cytokine expression in biliary cells (Figure 4). These bacteria did not have the same effect in hepatic cells, with the exception of *V. parvula*. These results are consistent with PSC disease progression, in which inflammation in the biliary tree precedes liver damage.^44^ Interestingly, *E. faecalis*, the primary strain that induced intestinal permeability, did not induce biliary inflammation according to our transcriptional analyses. These findings suggests that different PSC-associated bacteria may be responsible for unique aspects of cellular disease phenotypes. Additionally, while *E. coli, F. necrophorum*, and certain strains of *K. pneumoniae* were all H_2_S producers, we did not observe a stronger induction of inflammation by H_2_S-producing *K. pneumoniae* compared to non-producing *K. pneumoniae* (Table 2, Figure 4). Therefore, although bacterial H_2_S production may be associated with inflammation, as indicated by our results and as suggested by other studies,^45^ H_2_S production does not appear to be a primary driver of cytokine induction. Notably, reference strain *E. gallinarum* did not induce inflammatory cytokine expression in the tested biliary and hepatic human cancer cell lines but has been shown to induce inflammatory cytokines in murine hepatic cells derived from mice with a genetic predisposition to lupus-like autoimmunity.^21^ This result highlights the importance of cell type selection for modeling disease.

Our results identify TNFα and IL-8 as the primary cytokines induced by the PSC-associated bacteria (Figure 4). However, the other cytokines measured (IL-6, IL-17A, and IFNγ) have all been previously identified as elevated in PSC patients.^46,47^ A limitation of our cell assays is that only epithelial cells are included, not immune cells. It is likely that the presence of immune cells would result in the induction of more inflammatory cytokines by the PSC-associated bacteria. Nevertheless, our epithelial cell assays suggest that TNFα and IL-8 may be contributing to biliary inflammation at the local level. These cytokines have been previously shown to be elevated in PSC patient serum, with IL-8 also being significantly elevated in bile, consistent with our results.^48^

We observed that *S. salivarius, E. faecalis*, and *V. parvula* upregulate the expression of selected genes involved in sulfur metabolism, mucin expression, and the bicarbonate umbrella, perhaps in response to toxic factors produced by these bacteria. Notably, the bacteria affecting these protective pathways are not the same as the bacteria shown to induce inflammatory cytokines, with the exception of *V. parvula*. This finding again supports the notion that bacteria cultured from PSC patient bile affect different aspects of cellular disease phenotypes, with the exception of *V. parvula*, which appears to have a mild effect on all observed cellular responses. The two bacterial isolates that consistently produced H_2_S in vitro (*E. coli* and *F. necrophorum*) did not affect the expression of sulfur metabolism genes in host cells in these targeted analyses (Figure 5, Table 2). Thus, our results do not suggest that bacterial H_2_S production is driving expression of target sulfur metabolism genes. However, of potential relevance was our observation that supernatants from *V. dispar/nakazawae* cultures caused a decrease in mRNA expression of sulfide quinone oxidoreductase (SQOR) in biliary cells, which could lead to an impairment in H_2_S metabolism. Future work exploring the transcriptional and protein-level effects of H_2_S-producing bacteria on sulfur metabolism in host cells over time is necessary to uncover potential interactions between H_2_S production and host cell damage.

Overall, through the culturing of patient-derived bacterial isolates, the collection of sterile-filtered bacterial supernatant, and the use of human cellular assays under anaerobic conditions, we have demonstrated that factors produced by individual PSC-derived bacteria strains can induce disease-associated phenotypes in cells, including intestinal permeability, inflammation, and cell death. These results indicate that there may be a causal link between the presence of these bacteria in PSC patient microbiomes and disease development. Our work suggests a potential model for PSC pathogenesis in which increased intestinal permeability caused by *Enterococcus* or factors associated with co-occurrent IBD may allow bacteria to translocate from the gut to the liver where these microbes can colonize the biliary tree. Translocated bacteria such as *Escherichia, Fusobacterium*, and *Klebsiella* may then drive inflammation in bile ducts. This inflammation along with the prolonged exposure to cytotoxic factors produced by PSC-associated bacteria may then cause or contribute to the progression of biliary fibrosis and hepatic cirrhosis (Figure 6). Additionally, this model highlights the potential importance of the gut-liver axis in PSC,^2^ where dysbiosis in the gut microbiome and increased intestinal permeability may allow for both chronic disease and recurrence after liver transplant. These bacterial factors may contribute centrally to these critical steps in the pathogenesis of PSC, but likely operate in conjunction with other genetic, environmental or autoimmune contributions to intestinal permeability, inflammation, and cell death.

**Figure 6.**
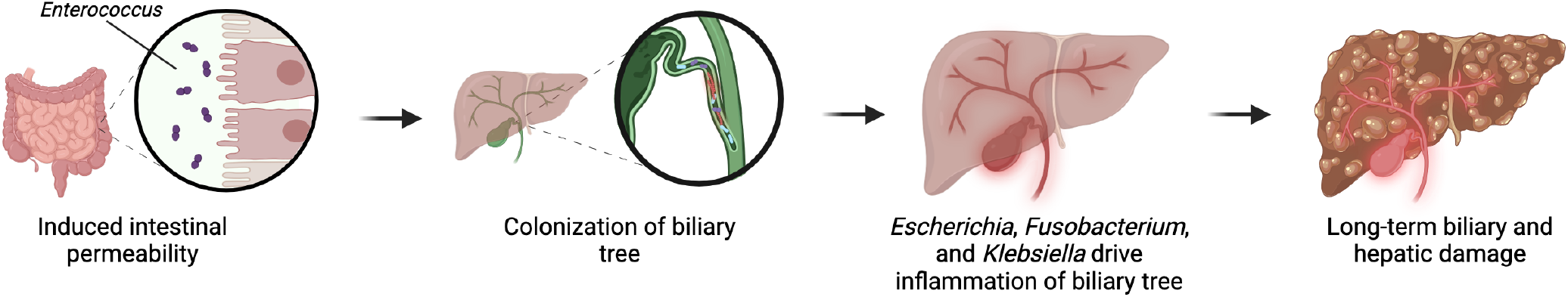
A potential model for microbial involvement in PSC pathogenesis.

The connections presented in this study between specific bacteria and disease-associated cellular responses may open up avenues for future therapeutic development. While our use of bacterial supernatants and biological replicates has allowed us to demonstrate a preliminary link between strains and cellular responses, future studies are needed to identify what specific factors, such as proteins or metabolites, are being produced by these bacteria and what quantity of these factors is needed to induce disease. This further examination could potentially allow for the identification of novel therapeutic targets for PSC.

## List of Abbreviations

CD: Crohn’s disease
ERCP: endoscopic retrograde cholangiography
IBD: inflammatory bowel disease
IC: indeterminate colitis
PSC: primary sclerosing cholangitis
UC: ulcerative colitis

## ACKNOWLEDGEMENTS

We would like to thank Martin Kriegel (Yale School of Medicine) for gifting *Enterococcus gallinarum*. Additionally, BioRender was used to create Figure 6 and the Graphical Abstract.

