## Supplemental Information for "Cultured Bacteria Isolated from Primary Sclerosing Cholangitis Patient Bile Induce Inflammation and Cell Death"

**Table S1.** Metagenomic sequencing statistics. NCBI BioProject database: PRJNA1169933.

| PSC Sample | Total Read Pairs | Total Reads (R1 + R2) | Total bp > Q30 | % bp > Q30 |
|---|---|---|---|---|
| PSC 1 | 5.93E+07 | 1.19E+08 | 1.53E+10 | 93.973 |
| PSC 1 – 8 months later | 3.31E+07 | 6.63E+07 | 9.03E+09 | 91.553 |
| PSC 2 | 3.68E+07 | 7.37E+07 | 1.01E+10 | 92.503 |
| PSC 3 | 6.17E+07 | 1.23E+08 | 1.59E+10 | 93.886 |
| PSC 4 | 3.88E+07 | 7.75E+07 | 1.07E+10 | 93.44 |
| PSC 6 | 3.66E+07 | 7.32E+07 | 1.01E+10 | 92.335 |
| PSC 7 | 2.76E+07 | 5.52E+07 | 7.51E+09 | 92.278 |
| PSC 9 | 2.77E+07 | 5.53E+07 | 7.65E+09 | 92.756 |

**Table S2.** Metagenomic data NCBI BioProject database (PRJNA1169933) accession numbers for each sample.

| | Accession # |
|---|---|
| **PSC 1** | SAMN44088305 |
| **PSC 2** | SAMN44088306 |
| **PSC 3** | SAMN44088307 |
| **PSC 4** | SAMN44088308 |
| **PSC 6** | SAMN44088309 |
| **PSC 7** | SAMN44088310 |
| **PSC 9** | SAMN44088311 |
| **PSC 1 (8 months later)** | SAMN44088312 |

**Table S3.** qPCR primers for human targets. Primer efficiencies were determined to be between 90 – 110% with $R^2$ values between 0.99 and 1.00.

| | PrimerBank ID | Forward | Reverse |
|---|---|---|---|
| **Muc1** | 324120973c1 | TGCCGCCGAAAGAACTACG | TGGGGTACTCGCTCATAGGAT |
| **Muc5AC** | 3334747a2 | CCATTGCTATTATGCCCTGTGT | TGGTGGACGGACAGTCACT |
| **TNF-alpha** | 25952110c2 | GAGGCCAAGCCCTGGTATG | CGGGCCGATTGATCTCAGC |
| **IL-6** | 224831235c1 | ACTCACCTCTTCAGAACGAATTG | CCATCTTTGGAAGGTTCAGGTTG |
| **IL-8** | 10834978a2 | ACTGAGAGTGATTGAGAGTGGAC | AACCCTCTGCACCCAGTTTTC |
| **IL-17A** | 27477085c1 | TCCCACGAAATCCAGGATGC | GGATGTTCAGGTTGACCATCAC |
| **IFNg** | 56786137c1 | TCGGTAACTGACTTGAATGTCCA | TCGCTTCCCTGTTTTAGCTGC |
| **AE2** | 314122223c1 | TCCTCCCACCACATCCATCA | CTCCTCAATGGTCGGGGTTTC |
| **SUOX** | 74099701c2 | ACTCAAGTCAATCCCCTCAAGG | GCTGGAGTTATCACCAGAGAAGG |
| **TST** | 34335291c2 | GACTGGACTCGGGCCATATC | ACGTGGCAATGAGAGGCTG |
| **SQOR** | 52851410c1 | AGCTAGAGTGACTGAGTTGAACC | AGCTGGATTCCGAGAGCAATAA |
| **UBC** | 305632811c1 | CTGGAAGATGGTCGTACCCTG | GGTCTTGCCAGTGAGTGTCT |

**Table S4.** Individual patient characteristics at time of bile collection

| Patient | Age | Sex | PSC Diagnosis | PSC Duration (years) | IBD Diagnosis | IBD Duration (years) | ERCP (#) | ALP (U/L) |
|---|---|---|---|---|---|---|---|---|
| PSC 1 | 38 | Male | PSC | 0 | IC | 22 | 1 | 58 |
| PSC 2 | 34 | Male | PSC large duct | 6 | UC/CD | n/a | 3 | 621 |
| PSC 3 | 74 | Male | PSC | 28 | UC | 52 | 6+ | 289 |
| PSC 4 | 67 | Female | PSC | 21 | None | | 6 | 143 |
| PSC 5 | 32 | Male | PSC | 8 | CD | 3 | 4 | 629 |
| PSC 6 | 38 | Male | PSC | 8 | UC/CD | 13 | 4 | 136 |
| PSC 7 | 28 | Male | PSC | 5 | None | | 1 | 554 |
| PSC 8 | 58 | Male | PSC | 10 | UC | 10+ | 4 | 472 |
| PSC 9 | 73 | Male | PSC | 3 | UC | 55 | 5 | 620 |
| PSC 10 | 41 | Female | PSC | 13 | IC | 14 | 8 | 207 |
| Cholecystectomy 1 | 59 | Female | None / Control | | | | 0 | 86 |
| Cholecystectomy 2 | 41 | Female | None / Control | | | | 0 | 41 |
| Cholecystectomy 3 | 35 | Male | None / Control | | | | 0 | 106 |

**Table S5.** PSC 1 Colony 16S Sanger Sequencing and $H_2S$ Production

| Colony Number | BLAST Result | $H_2S$ Production? |
|---|---|---|
| 1 | *Streptococcus salivarius* | N |
| 2 | *Streptococcus salivarius* | N |
| 3 | *Streptococcus salivarius* | N |
| 4 | *Streptococcus salivarius* | N |
| 5 | *Streptococcus salivarius* | N |
| 6 | *Streptococcus salivarius* | N |
| 7 | No priming | N |
| 8 | No priming | N |
| 9 | *Streptococcus salivarius* | N |
| 10 | *Streptococcus salivarius* | N |
| 11 | *Streptococcus salivarius* | N |
| 12 | *Streptococcus salivarius* | N |
| 13 | *Streptococcus salivarius* | N |
| 14 | *Streptococcus salivarius* | N |
| 15 | *Streptococcus salivarius* | N |
| 16 | *Streptococcus salivarius* | N |
| 17 | *Streptococcus salivarius* | N |
| 18 | *Paraclostridium bifermentans/benzoelyticum* | Y |
| 19 | *Streptococcus salivarius* | N |
| 20 | *Streptococcus salivarius* | N |
| 21 | *Streptococcus salivarius* | N |
| 22 | No priming | N |
| 23 | *Streptococcus salivarius* | N |
| 24 | *Streptococcus salivarius* | N |
| 25 | *Streptococcus salivarius* | N |
| 26 | *Streptococcus salivarius* | N |
| 27 | *Streptococcus salivarius* | N |
| 28 | *Streptococcus salivarius* | N |
| 29 | *Streptococcus salivarius* | N |
| 30 | *Streptococcus salivarius* | N |
| 31 | *Streptococcus salivarius* | N |
| 32 | *Streptococcus salivarius* | N |
| 33 | *Streptococcus salivarius* | N |
| 34 | *Streptococcus salivarius* | N |
| 35 | *Streptococcus salivarius* | N |
| 36 | *Streptococcus salivarius* | N |
| 37 | *Streptococcus salivarius* (Early termination) | N |
| 38 | *Streptococcus salivarius* | N |
| 39 | *Streptococcus salivarius* | N |
| 40 | *Streptococcus salivarius* | N |
| 41 | *Streptococcus salivarius* | N |
| 42 | *Streptococcus salivarius* | N |
| 43 | *Streptococcus salivarius* | N |
| 44 | *Streptococcus salivarius* | N |
| 45 | *Streptococcus salivarius* | N |
| 46 | *Streptococcus salivarius* | N |
| 47 | *Streptococcus salivarius* | N |
| 48 | *Streptococcus salivarius* | N |
| 49 | *Streptococcus salivarius* | N |
| 50 | *Streptococcus salivarius* | N |

**Table S6.** PSC 2 Colony 16S Sanger Sequencing and $H_2S$ Production

| Colony Number | BLAST Result | $H_2S$ Production? |
|---|---|---|
| 1 | *Streptococcus salivarius* | N |
| 2 | *Streptococcus salivarius* | N |
| 3 | *Streptococcus salivarius* | N |
| 4 | *Streptococcus salivarius* | N |
| 5 | *Streptococcus sp. / Streptococcus anginosus* | N |
| 6 | *Streptococcus salivarius* | N |
| *7* | *Streptococcus salivarius* | N |
| *8* | *Streptococcus salivarius* | N |
| 9 | *Streptococcus salivarius* | N |
| 10 | *Streptococcus salivarius* | N |
| 11 | *Streptococcus sp. / Streptococcus anginosus* | N |
| 12 | *Streptococcus sp. / Streptococcus anginosus* | N |
| 13 | *Streptococcus salivarius* | N |
| 14 | *Streptococcus salivarius* | N |
| 15 | *Streptococcus salivarius* | N |
| 16 | *Streptococcus salivarius* | N |
| 17 | *Streptococcus salivarius* | N |
| 18 | *Streptococcus salivarius* | N |
| 19 | *Streptococcus salivarius* | N |
| 20 | *Streptococcus salivarius* | N |
| 21 | *Streptococcus salivarius* | N |
| *22* | *Streptococcus salivarius* | N |
| 23 | *Streptococcus salivarius* | N |
| 24 | *Streptococcus sp. / Streptococcus anginosus* | N |
| 25 | No priming | N |
| 26 | *Streptococcus salivarius* | N |
| 27 | *Streptococcus salivarius* | N |
| 28 | *Streptococcus salivarius* | N |
| 29 | *Streptococcus salivarius* | N |
| 30 | *Streptococcus salivarius* | N |
| 31 | *Streptococcus salivarius* | N |
| 32 | *Streptococcus salivarius* | N |
| 33 | *Streptococcus salivarius* | N |
| 34 | *Streptococcus salivarius* | N |
| 35 | *Streptococcus salivarius* | N |
| 36 | *Streptococcus salivarius* | N |
| 37 | *Streptococcus salivarius* | N |
| 38 | *Streptococcus salivarius* | N |
| 39 | No priming | N |
| 40 | *Streptococcus salivarius* | N |
| 41 | *Streptococcus salivarius* | N |
| 42 | *Streptococcus salivarius* | N |
| 43 | *Streptococcus salivarius* | N |
| 44 | *Streptococcus salivarius* | N |
| 45 | *Streptococcus salivarius* | N |
| 46 | *Streptococcus salivarius* | N |
| 47 | *Streptococcus salivarius* | N |
| 48 | No priming | N |
| 49 | *Streptococcus salivarius* | N |
| 50 | *Streptococcus salivarius* | N |

**Table S7.** PSC 3 Colony 16S Sanger Sequencing and $H_2S$ Production

| Colony Number | BLAST Result | $H_2S$ Production? |
|---|---|---|
| 1 | *Klebsiella pneumoniae* | N |
| 2 | *Klebsiella pneumoniae* | N |
| 3 | *Klebsiella pneumoniae* | N |
| 4 | *Klebsiella pneumoniae* | N |
| 5 | *Klebsiella pneumoniae* | N |
| 6 | *Klebsiella pneumoniae* | N |
| 7 | *Klebsiella pneumoniae* | N |
| 8 | *Klebsiella pneumoniae* | N |
| 9 | *Klebsiella pneumoniae* | N |
| 10 | *Klebsiella pneumoniae* | N |
| 11 | *Klebsiella pneumoniae* | N |
| 12 | *Klebsiella pneumoniae* | N |
| 13 | *Klebsiella pneumoniae* | N |
| 14 | *Klebsiella pneumoniae* | N |
| 15 | *Klebsiella pneumoniae* | N |
| 16 | *Klebsiella pneumoniae* | N |
| 17 | *Klebsiella pneumoniae* | N |
| 18 | *Klebsiella pneumoniae* | N |
| 19 | *Klebsiella pneumoniae* | N |
| 20 | *Klebsiella pneumoniae* | N |
| 21 | *Klebsiella pneumoniae* | N |
| 22 | *Klebsiella pneumoniae* | N |
| 23 | *Klebsiella pneumoniae* | N |
| 24 | *Klebsiella pneumoniae* | N |
| 25 | *Klebsiella pneumoniae* | N |
| 26 | *Klebsiella pneumoniae* | N |
| 27 | *Klebsiella pneumoniae* | N |
| 28 | *Klebsiella pneumoniae* | N |
| 29 | *Klebsiella pneumoniae* | N |
| 30 | *Klebsiella pneumoniae* | N |
| 31 | *Klebsiella pneumoniae* | N |
| 32 | *Klebsiella pneumoniae* | N |
| 33 | *Klebsiella pneumoniae* | N |
| 34 | *Klebsiella pneumoniae* | N |
| 35 | *Klebsiella pneumoniae* | N |
| 36 | *Klebsiella pneumoniae* | N |
| 37 | *Klebsiella pneumoniae* | N |
| 38 | *Klebsiella pneumoniae* | N |
| 39 | *Klebsiella pneumoniae* | N |
| 40 | *Klebsiella pneumoniae* | N |
| 41 | *Klebsiella pneumoniae* | N |
| 42 | *Klebsiella pneumoniae* | N |
| 43 | *Klebsiella pneumoniae* | N |
| 44 | *Klebsiella pneumoniae* | N |
| 45 | *Klebsiella pneumoniae* | N |
| 46 | *Klebsiella pneumoniae* | N |
| 47 | *Klebsiella pneumoniae* | N |
| 48 | *Klebsiella pneumoniae* | N |
| 49 | *Klebsiella pneumoniae* | N |
| 50 | *Klebsiella pneumoniae* | N |

**Table S8.** PSC 6 Colony 16S Sanger Sequencing and $H_2S$ Production

| Colony Number | BLAST Result | $H_2S$ Production? |
|---|---|---|
| 1 | *Enterococcus faecalis* | N |
| 2 | *Enterococcus faecalis* | N |
| 3 | *Enterococcus faecalis* | N |
| 4 | *Enterococcus faecalis* | N |
| 5 | *Enterococcus faecalis* | N |
| 6 | *Enterococcus faecalis* | N |
| 7 | *Enterococcus faecalis* | N |
| 8 | *Enterococcus faecalis* | N |
| 9 | *Enterococcus faecalis* | N |
| 10 | *Enterococcus faecalis* | N |
| 11 | *Enterococcus faecalis* | N |
| 12 | *Enterococcus faecalis* | N |
| 13 | *Enterococcus faecalis* | N |
| 14 | *Enterococcus faecalis* | N |
| 15 | *Enterococcus faecalis* | N |
| 16 | *Enterococcus faecalis* | N |
| 17 | *Enterococcus faecalis* | N |
| 18 | *Enterococcus faecalis* | N |
| 19 | *Enterococcus faecalis* | N |
| 20 | *Enterococcus faecalis* | N |
| 21 | *Enterococcus faecalis* | N |
| 22 | *Enterococcus faecalis* | N |
| 23 | *Enterococcus faecalis* | N |
| 24 | *Enterococcus faecalis* | N |
| 25 | *Enterococcus faecalis* | N |
| 26 | *Enterococcus faecalis* | N |
| 27 | No priming | N |
| 28 | No priming | N |
| 29 | *Enterococcus faecalis* | N |
| 30 | *Enterococcus faecalis* | N |
| 31 | *Enterococcus faecalis* | N |
| 32 | *Enterococcus faecalis* | N |
| 33 | *Enterococcus faecalis* | N |
| 34 | *Enterococcus faecalis* | N |
| 35 | *Enterococcus faecalis* | N |
| 36 | *Enterococcus faecalis* | N |
| 37 | *Enterococcus faecalis* | N |
| 38 | *Enterococcus faecalis* | N |
| 39 | *Enterococcus faecalis* | N |
| 40 | *Enterococcus faecalis* | N |
| 41 | *Enterococcus faecalis* | N |
| 42 | No priming | N |
| 43 | *Enterococcus faecalis* | N |
| 44 | *Enterococcus faecalis* | N |
| 45 | *Enterococcus faecalis* | N |
| 46 | *Enterococcus faecalis* | N |
| 47 | *Enterococcus faecalis* | N |
| 48 | *Enterococcus faecalis* | N |
| 49 | *Enterococcus faecalis* | N |
| 50 | No priming | N |

**Table S9.** PSC 7 Colony 16S Sanger Sequencing and $H_2S$ Production

| Colony Number | BLAST Result | $H_2S$ Production? |
|---|---|---|
| 1 | *Schaalia odontolytica / Actinomyces sp.* | N |
| 2 | *Streptococcus parasanguinis/S. australis* | N |
| 3 | *Prevotella melaninogenica* | N |
| 4 | *Actinomyces sp. / Schaalia odontolytica* | N |
| 5 | *Veillonella parvula* | N |
| 6 | *Campylobacter showae* | N |
| 7 | *Streptococcus sp. / Uncultured organism clone* | N |
| 8 | *Actinomyces sp. / Schaalia odontolytica* | N |
| 9 | *Streptococcus salivarius* | N |
| 10 | *Actinomyces sp. / Schaalia odontolytica* | N |
| 11 | *Campylobacter concisus* | N |
| 12 | *Streptococcus salivarius / Uncultured bacterium clone* | N |
| 13 | *Actinomyces oris / Uncultured bacterium clone* | N |
| 14 | *Veillonella nakazawae / V. dispar* | N |
| 15 | *Veillonella nakazawae / V. dispar* | N |
| 16 | *Schaalia odontolyica* | N |
| 17 | *Streptococcus anginosus* | N |
| 18 | *Actinomyces sp. / Schaalia odontolytica* | N |
| 19 | *Actinomyces sp. / Schaalia odontolytica* | N |
| 20 | *Streptococcus pseudopneumoniae/ S. symci / S. mitis / S. pneumoniae* | N |
| 21 | *Streptococcus sp. / Streptococcus mitis* | N |
| 22 | *Schaalia odontolytica / Actinomyces sp.* | N |
| 23 | *Schaalia odontolyica / Actinomyces odontolyticus* | N |
| 24 | *Uncultured bacterium / Veillonella nakazawae / V. dispar* | N |
| 25 | *Schaalia odontolytica / Actinomyces sp.* | N |
| 26 | *Actinomyces sp. / Schaalia odontolytica* | N |
| 27 | *Streptococcus parasanguinis / S. australis* | N |
| 28 | No priming | N |
| 29 | *Streptococcus oralis* | N |
| 30 | *Streptococcus parasanguinis / S. australis* | N |
| 31 | *Streptococcus pseudopneumoniae / S. oralis / S. mitis* | N |
| 32 | *Schaalia odontolytica / Actinomyces sp.* | N |
| 33 | *Gemella sp. / G. haemolysans / Uncultured bacterium clone* | N |
| 34 | *Streptococcus pseudopneumoniae / S. oralis / S. mitis* | N |
| 35 | *Streptococcus pseudopneumoniae / S. oralis / S. mitis* | N |
| 36 | *Lachnoanaerobaculum sp.* | N |
| 37 | *Streptococcus pseudopneumoniae / S. symci / S. mitis / S. pneumoniae* | N |
| 38 | *Veillonella nakazawae / V. dispar* | N |
| 39 | *Uncultured organism clone / Streptococcus sp.* | N |
| 40 | *Streptococcus salivarius* | N |
| 41 | *Actinomyces sp. / Schaalia odontolytica* | N |
| 42 | *Streptococcus pseudopneumoniae / S. symci / S. mitis / S. pneumoniae* | N |
| 43 | *Actinomyces sp. / Schaalia odontolytica* | N |
| 44 | *Streptococcus australis / S. infantis / Uncultured organism clone* | N |
| 45 | *Streptococcus pseudopneumoniae / S. symci / S. mitis / S. pneumoniae* | N |
| 46 | *Prevotella melaninogenica* | N |
| 47 | *Actinomyces sp. / Schaalia odontolytica* | N |
| 48 | *Veillonella nakazawae / V. dispar* | N |
| 49 | *Streptococcus salivarius* | N |
| 50 | *Streptococcus infantis / S. australis* | N |

**Table S10.** PSC 8 Colony 16S Sanger Sequencing and $H_2S$ Production

| Colony Number | BLAST Result | $H_2S$ Production? |
|---|---|---|
| 1 | No Priming | N |
| 2 | *Fusobacterium necrophorum* | Y |
| 3 | *Klebsiella pneumoniae/variicola (non-specific)* | Y |
| 4 | *Fusobacterium necrophorum* | N |
| 5 | *Streptococcus anginosus* | N |
| 6 | *Klebsiella pneumoniae/variicola* | N |
| 7 | *Klebsiella pneumoniae/variicola* | Y |
| 8 | *Fusobacterium necrophorum* | Y |
| 9 | *Klebsiella pneumoniae/variicola* | N |
| 10 | *Fusobacterium necrophorum* | Y |
| 11 | *Fusobacterium necrophorum* | Y |
| 12 | *Klebsiella pneumoniae/variicola* | N |
| 13 | *Fusobacterium necrophorum* | Y |
| 14 | *Fusobacterium necrophorum* | N |
| 15 | *Klebsiella pneumoniae/variicola* | N |
| 16 | *Klebsiella pneumoniae/variicola* | N |
| 17 | *Campylobacter gracilis* | N |
| 18 | *Fusobacterium necrophorum* | Y |
| 19 | No Priming | N |
| 20 | *Klebsiella pneumoniae/variicola* | N |
| 21 | *Fusobacterium necrophorum* | Y |
| 22 | *Fusobacterium necrophorum* | Y |
| 23 | *Fusobacterium necrophorum* | Y |
| 24 | *Fusobacterium necrophorum* | Y |
| 25 | *Fusobacterium necrophorum* | Y |
| 26 | *Fusobacterium necrophorum* | Y |
| 27 | *Klebsiella pneumoniae/variicola* | N |
| 28 | *Klebsiella pneumoniae/variicola* | N |
| 29 | *Klebsiella pneumoniae/variicola* | N |
| 30 | *Klebsiella pneumoniae/variicola* | N |
| 31 | *Fusobacterium necrophorum* | Y |
| 32 | *Klebsiella pneumoniae/variicola* | N |
| 33 | *Klebsiella pneumoniae/variicola* | N |
| 34 | *Fusobacterium necrophorum* | Y |
| 35 | *Fusobacterium necrophorum* | Y |
| 36 | *Fusobacterium necrophorum* | Y |
| 37 | *Klebsiella pneumoniae/variicola* | N |
| 38 | *Fusobacterium necrophorum* | Y |
| 39 | *Klebsiella pneumoniae/variicola* | N |
| 40 | *Klebsiella pneumoniae/variicola* | N |
| 41 | *Fusobacterium necrophorum* | Y |
| 42 | *Klebsiella pneumoniae/variicola* | N |
| 43 | *Fusobacterium necrophorum* | Y |
| 44 | *Prevotella phocaeensis/oralis* | Y |
| 45 | *Fusobacterium necrophorum* | Y |
| 46 | *Klebsiella pneumoniae/variicola* | N |
| 47 | *Fusobacterium necrophorum* | Y |
| 48 | *Klebsiella pneumoniae/variicola* | N |
| 49 | *Fusobacterium necrophorum* | Y |
| 50 | *Fusobacterium necrophorum* | Y |

**Table S11.** PSC 9 Colony 16S Sanger Sequencing and $H_2S$ Production

| Colony Number | BLAST Result | $H_2S$ Production? |
|---|---|---|
| 1 | *Streptococcus anginosus* | N |
| 2 | *Klebsiella pneumoniae* | N |
| 3 | *Streptococcus anginosus* | N |
| 4 | *Klebsiella pneumoniae* | N |
| 5 | *Streptococcus anginosus* | N |
| 6 | *Klebsiella pneumoniae* | N |
| 7 | *Enterococcus casseliflavus / gallinarum* | N |
| 8 | *Fusobacterium animalis* | Y |
| 9 | *Enterococcus casseliflavus / gallinarum* | N |
| 10 | *Streptococcus anginosus* | N |
| 11 | *Enterococcus casseliflavus* | N |
| 12 | *Streptococcus anginosus* | N |
| 13 | *Clostridium perfringens* | Y |
| 14 | *Clostridium perfringens* | Y |
| 15 | *Klebsiella pneumoniae* | N |
| 16 | *Streptococcus anginosus* | N |
| 17 | *Klebsiella pneumoniae* | N |
| 18 | *Klebsiella pneumoniae / quasipneumoniae* | N |
| 19 | *Streptococcus anginosus* | N |
| 20 | *Klebsiella pneumoniae* | N |
| 21 | *Streptococcus anginosus* | N |
| 22 | *Klebsiella pneumoniae / quasipneumoniae* | N |
| 23 | *Klebsiella pneumoniae* | N |
| 24 | *Streptococcus anginosus* | N |
| 25 | *Klebsiella pneumoniae* | N |
| 26 | *Klebsiella pneumoniae* | Y |
| 27 | *Klebsiella pneumoniae / quasipneumoniae* | N |
| 28 | *Klebsiella pneumoniae / quasipneumoniae* | N |
| 29 | *Streptococcus anginosus* | N |
| 30 | *Klebsiella pneumoniae* | N |
| 31 | *Streptococcus anginosus* | N |
| 32 | *Klebsiella pneumoniae* | N |
| 33 | *Klebsiella pneumoniae* | N |
| 34 | *Klebsiella pneumoniae* | N |
| 35 | *Streptococcus anginosus* | N |
| 36 | *Streptococcus anginosus* | N |
| 37 | *Klebsiella pneumoniae* | N |
| 38 | *Enterococcus casseliflavus* | N |
| 39 | *Enterococcus casseliflavus* | N |
| 40 | *Enterococcus casseliflavus / gallinarum* | N |
| 41 | *Streptococcus anginosus* | N |
| 42 | *Enterococcus casseliflavus / gallinarum* | N |
| 43 | *Klebsiella pneumoniae* | N |
| 44 | *Streptococcus anginosus* | Y |
| 45 | *Streptococcus anginosus* | N |
| 46 | *Klebsiella pneumoniae* | N |
| 47 | *Streptococcus anginosus* | N |
| 48 | *Klebsiella pneumoniae* | N |
| 49 | *Klebsiella pneumoniae* | Y |
| 50 | *Streptococcus anginosus* | N |

**Table S12.** PSC 10 Colony 16S Sanger Sequencing and $H_2S$ Production

| Colony Number | BLAST Result | $H_2S$ Production? |
|---|---|---|
| 1 | *Escherichia coli* | Y |
| 2 | *Escherichia coli* | Y |
| 3 | *Escherichia coli* | Y |
| 4 | *Escherichia coli* | Y |
| 5 | *Escherichia coli* | Y |
| 6 | *Escherichia coli* | Y |
| 7 | *Escherichia coli* | Y |
| 8 | *Escherichia coli* | Y |
| 9 | *Escherichia coli* | Y |
| 10 | *Escherichia coli* | Y |
| 11 | *Escherichia coli* | Y |
| 12 | *Escherichia coli* | Y |
| 13 | *Escherichia coli* | Y |
| 14 | *Escherichia coli* | Y |
| 15 | *Shigella sonnei/Eschericha coli* | Y |
| 16 | *Escherichia coli* | Y |
| 17 | *Shigella sonnei/Eschericha coli* | Y |
| 18 | *Shigella sonnei/Eschericha coli* | Y |
| 19 | *Escherichia coli* | Y |
| 20 | *Escherichia coli* | Y |
| 21 | *Escherichia coli* | Y |
| 22 | *Escherichia coli* | Y |
| 23 | *Escherichia coli* | Y |
| 24 | *Shigella sonnei/Eschericha coli* | Y |
| 25 | *Escherichia coli* | Y |
| 26 | *Escherichia marmotae/ coli* | Y |
| 27 | *Escherichia coli* | Y |
| 28 | *Escherichia coli* | Y |
| 29 | *Shigella sonnei/Eschericha coli* | Y |
| 30 | *Shigella sonnei/Eschericha coli* | Y |
| 31 | *Shigella sonnei/Eschericha coli* | Y |
| 32 | *Escherichia coli* | Y |
| 33 | *Escherichia coli* | Y |
| 34 | *Escherichia coli* | Y |
| 35 | *Escherichia coli* | Y |
| 36 | *Escherichia coli* | Y |
| 37 | *Escherichia coli* | Y |
| 38 | *Shigella sonnei/Eschericha coli* | Y |
| 39 | *Escherichia coli* | Y |
| 40 | *Escherichia coli* | Y |
| 41 | *Shigella sonnei/Eschericha coli* | Y |
| 42 | *Shigella sonnei/Eschericha coli* | Y |
| 43 | *Shigella sonnei/Eschericha coli* | Y |
| 44 | *Escherichia coli* | Y |
| 45 | *Shigella sonnei/Eschericha coli* | Y |
| 46 | *Escherichia coli* | Y |
| 47 | *Escherichia coli* | Y |
| 48 | *Escherichia coli* | Y |
| 49 | *Escherichia marmotae/ coli* | Y |
| 50 | *Escherichia coli* | Y |

**Table S13.** PSC 1 (8 Months Later) Colony 16S Sanger Sequencing and $H_2S$ Production

| Colony Number | BLAST Result | $H_2S$ Production? |
|---|---|---|
| 1 | *Streptococcus salivarius* | N |
| 2 | *Streptococcus salivarius* | N |
| 3 | *Streptococcus salivarius* | N |
| 4 | *Neisseria perflava* | N |
| 5 | *Staphylococcus epidermidis/caprae* | N |
| *6* | No priming | *N* |
| 7 | *Streptococcus salivarius* | N |
| 8 | *Streptococcus salivarius* | N |
| 9 | *Campylobacter concisus* | N |
| 10 | *Streptococcus salivarius* | N |
| 11 | *Streptococcus salivarius* | N |
| 12 | *Streptococcus salivarius* | N |
| 13 | *Streptococcus salivarius* | N |
| 14 | *Streptococcus salivarius* | N |
| 15 | *Streptococcus salivarius* | N |
| 16 | *Schaalia odontolytica* (non-specific) | N |
| 17 | *Streptococcus salivarius* | N |
| 18 | *Streptococcus salivarius* | N |
| 19 | *Streptococcus parasanguinis* | N |
| 20 | Poor quality | N |
| 21 | No priming | N |
| 22 | *Streptococcus salivarius* | N |
| 23 | *Streptococcus salivarius* | N |
| 24 | *Streptococcus pneumoniae/australis/rubneri* | N |
| 25 | Poor quality | N |
| 26 | Poor quality | N |
| 27 | *Streptococcus salivarius* | N |
| 28 | No priming | N |
| 29 | No priming | N |
| 30 | *Streptococcus salivarius* | N |
| 31 | *Streptococcus salivarius* | N |
| 32 | *Streptococcus salivarius* | N |
| 33 | *Campylobacter concisus* | N |
| 34 | *Streptococcus salivarius* | N |
| 35 | *Streptococcus salivarius* | N |
| 36 | *Streptococcus salivarius* | N |
| 37 | *Streptococcus salivarius* | N |
| 38 | *Streptococcus salivarius* | N |
| 39 | *Streptococcus salivarius* | N |
| 40 | *Streptococcus salivarius* | N |
| 41 | *Streptococcus salivarius* (Poor quality) | N |
| 42 | *Streptococcus salivarius* | N |
| 43 | *Streptococcus salivarius* | N |
| 44 | *Streptococcus salivarius* | N |
| 45 | *Streptococcus salivarius* | N |
| 46 | *Streptococcus salivarius* | N |
| 47 | *Streptococcus salivarius* | N |
| 48 | *Streptococcus salivarius* | N |
| 49 | *Streptococcus salivarius* | N |
| 50 | Poor quality | N |

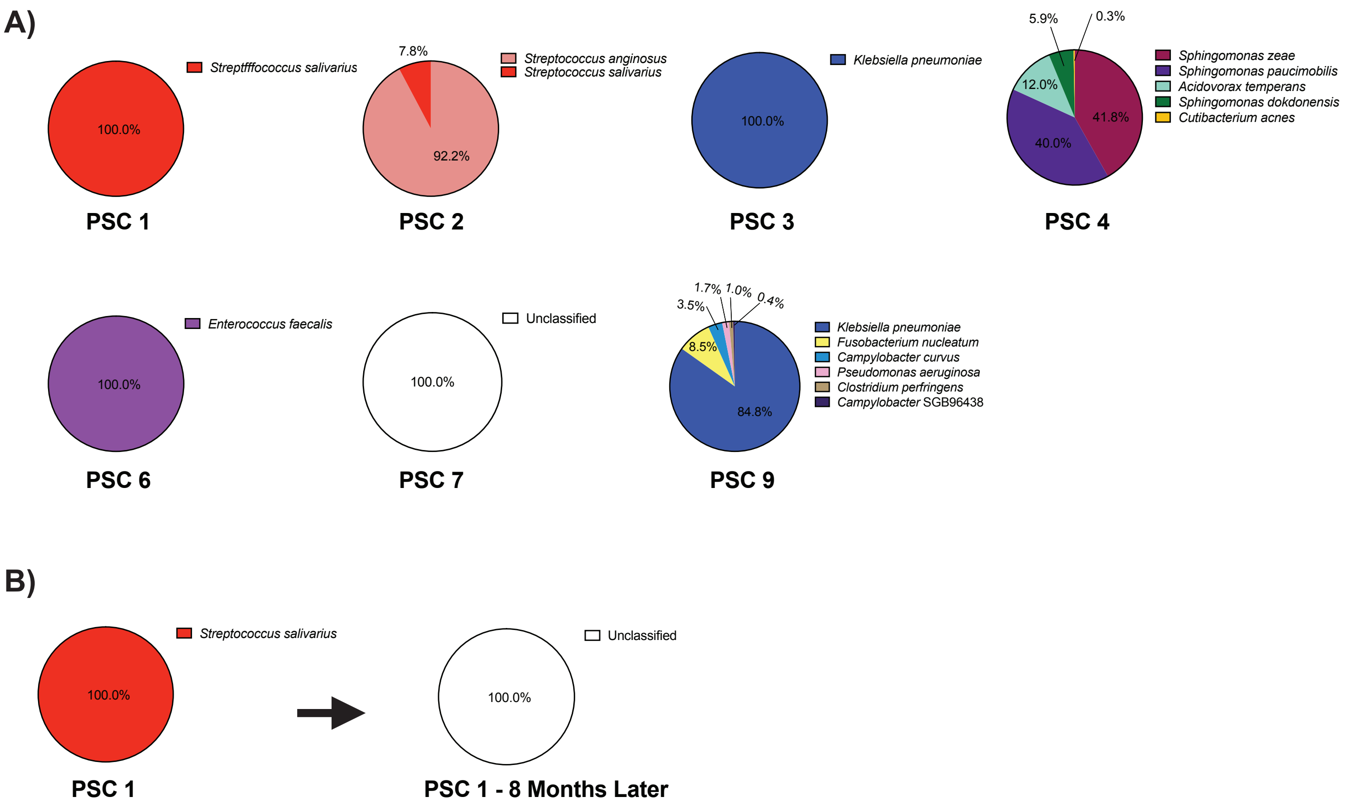


**Figure S1. Relative abundance of bacterial taxa in human bile samples as determined by shotgun metagenomic sequencing.** Each pie chart represents the microbial community composition of a distinct human bile sample. The relative abundances of each bacterial taxa are expressed as percentages of the total microbial community within each sample. Taxonomic profiling and relative abundances were determined using MetaPhlan 4.0.[1] NCBI BioProject database: PRJNA1169933.

**Table S14.** Source for bacteria used in cellular assays

| | Source |
|---|---|
| ***Enterococcus faecalis*** | PSC 6, Colony 1 |
| ***Escherichia coli*** | PSC 10, Colony 1 |
| ***Fusobacterium necrophorum*** | PSC 8, Colony 2 |
| ***Klebsiella pneumoniae* (H$_2$S producer)** | PSC 9, Colony 49 |
| ***Klebsiella pneumoniae* (non-producer)** | PSC 3, Colony 1 |
| ***Streptococcus salivarius*** | PSC 1, Colony 1 |
| ***Veillonella dispar/nakazawae*** | PSC 7, Colony 14 |
| ***Veillonella parvula*** | PSC 7, Colony 5 |

**Table S15.** Relative abundance profiling of metagenomic data using Kraken2[2] followed by KronaTools.[3] NCBI BioProject database: PRJNA1169933.

| | Bacteria (%) | Eukaryota (%) |
|---|---|---|
| **PSC 1** | 3 | 88 |
| **PSC 2** | 18 | 71 |
| **PSC 3** | 24 | 63 |
| **PSC 4** | 18 | 69 |
| **PSC 6** | 18 | 71 |
| **PSC 7** | 0.4 | 91 |
| **PSC 9** | 58 | 24 |
| **PSC 1 (8 months later)** | 1 | 91 |
